# Effects of high temperature and heavy precipitation on drinking water quality and child hand contamination levels in rural kenya

**DOI:** 10.1101/2022.10.04.510863

**Authors:** Julie E. Powers, Maryanne Mureithi, John Mboya, Jake Campolo, Jenna M. Swarthout, Joseph Pajka, Clair Null, Amy J. Pickering

## Abstract

Climate change may impact human health through the influence of weather on environmental transmission of diarrhea. Previous studies have found that high temperatures and heavy precipitation are associated with increased diarrhea prevalence, but the underlying causal mechanisms are not clear. We linked measurements of *Escherichia coli* in source water (n=1,673), stored drinking water (n=8,924), and hand rinses from children <2 years old (n=2,660) with publicly available gridded temperature and precipitation data (at ≤0.2 degree spatial resolution and daily temporal resolution) by the GPS coordinates and date of sample collection. Measurements were collected over a 3-year period across a 2500 km^2^ area in rural Kenya. In drinking water sources, high 7-day temperature was associated with a 0.16 increase in log_10_ *E. coli* levels (p<0.001), while heavy 7-day total precipitation was associated with a 0.29 increase in log_10_ *E. coli* levels (p<0.001). In household stored drinking water, heavy 7-day precipitation was associated with a 0.079 increase in log_10_ *E. coli* levels (p=0.042). On child hands, high 7-day temperature was associated with a 0.39 decrease in log_10_ *E. coli* levels (p<0.001). Our findings provide insight on how climate change could impact environmental transmission of bacterial pathogens in Kenya, and suggest water treatment could be a mitigation strategy.

## INTRODUCTION

In 2015, diarrhea was the ninth leading cause of death in all ages of people and the fourth leading cause of death in children under five^1^. Diarrhea also contributes to malnutrition^2–5^, stunting^6–12^, and cognitive impairment^6,7,13–20^ that could extend into adulthood^7,21^. Diarrhea is caused by enteric pathogen infections (bacterial, viral, or parasitic^22^) that are transmitted via the fecal-oral route: contaminated feces from an infected human or animal spread through environmental pathways and are ingested by another person^22–26^. In recent years, progress has been made in reducing the global burden of diarrhea: Among children under 5 years between 2005 and 2015, the mortality rate due to diarrhea (deaths due to diarrhea per population) decreased by 39.2% and diarrhea incidence decreased by 10.4%^1^. Diarrhea incidence has not decreased as quickly as diarrhea-associated mortality, suggesting that improved access to treatment may be largely responsible for the reductions in mortality. Extreme weather associated with climate change could threaten recent progress^27^ because temperature^28–34^ and heavy rainfall^31,34–36^ are positively associated with diarrhea. A systematic review by Levy et al. identified 53 quantitative analyses (65%) showing a significant positive association between temperature and diarrhea and 10 quantitative analyses (71%) showing a significant positive association between heavy rainfall events and diarrhea^37^. Some studies have found that the association between rainfall and diarrhea only holds under certain conditions, such as following prolonged dry periods^35,38,39^. Notably, of the 141 studies identified by Levy et al., only 11 (< 8%) were conducted in Sub-Saharan Africa^37,40^, suggesting a need for additional evidence in this geographic area.

The underlying causal mechanisms for the relationships between weather and diarrhea are not well established^41,42^. Heavy rainfall may cause surface runoff and flooding, potentially transporting feces and contaminating household environments and drinking water sources. However, heavy rainfall could also dilute the concentration of fecal matter in drinking water sources. High temperatures may influence pathogen survival in the environment, but the direction of the effect is unclear: pathogens may die off at a faster rate under high temperature conditions, but growth could also accelerate if sufficient nutrients are present^43,44^. A few recent studies found that heavy rainfall was associated with increased *Escherichia coli* (fecal indicator bacteria) levels in drinking water sources^40,45–47^ and household stored water^45,46^ in locations in Bangladesh^46,47^, Burkina Faso^40^, Nepal^46^, and Tanzania^45,46^. Higher temperatures increased *E. coli* levels in Bangladesh and Nepal, but decreased *E. coli* levels in Tanzania^46^. This variation by location suggests that the effects of weather on water quality are highly context-specific and underscores the need for evidence from additional locations.

In addition to physical and biological mechanisms, temperature and precipitation extremes could also lead to community or household-level behavioral changes that influence exposure to pathogens^46^. At the community level, agricultural activities such as application of animal feces as fertilizer may be correlated with temperature and precipitation. Planting occurs twice per year for many common crops in this region, once in February to March and again in August to October^48^. February to March is typically warmer than average, and both planting periods directly precede the rainy seasons, which occur from March to May and from October to December^49^. At the household level, it is common to use multiple water sources^50,51^. Households may choose to collect rainwater after periods of heavy rainfall. If heavy rainfall or high temperatures lead to perceived changes in water quality (e.g., color or turbidity), households may react to these changes and decide to switch sources or treat their water.

On a global level, as mean surface temperature rises, extreme precipitation events are projected to become more frequent and intense, and heat waves are projected to become more frequent and longer in duration^52^. In East Africa, mean annual temperature is expected to increase 2-4 degrees Celsius by 2050^53^. Precipitation projections vary widely: some models predict a potential increase of 2-4 extreme precipitation events annually in East Africa (Kenya and Tanzania)^54^ while others predict an increase in intensity and density of extreme precipitation events, but not a change in the actual number of events^55^. Kolstad & Johansson estimated that projected regional warming will increase the relative risk of diarrhea in equatorial Africa by 23% by the end of the 21^st^ century, threatening recent progress^27^. However, large uncertainties are associated with projecting the severity of the impact on diarrhea due to climate change, and more empirical data is needed to better understand the relationships between climate and human health^27^. The goal of this work is to examine associations between weather (heavy 7-day precipitation and high 7-day temperature) and environmental *E. coli* contamination (source water, stored water, and child hands) in Kenyan households. To our knowledge, this is the first study to examine the effects of weather on hand contamination.

## MATERIALS AND METHODS

### Data Sources

We leveraged environmental *E. coli* contamination data from the WASH Benefits Study in western Kenya (Figure 1), a multiyear randomized controlled trial that studied the effects of water, sanitation, hygiene, and nutrition interventions on diarrhea and growth in children during their first two years of life^56,57^. Investigators designed the trial with a control arm (C) and six intervention arms: water treatment (W); sanitation (S); handwashing with soap (H); combined water, sanitation, and handwashing (WSH); nutrition (N); and combined water, sanitation, handwashing, and nutrition (WSHN) (see Supplementary Methods).

**Figure 1:**
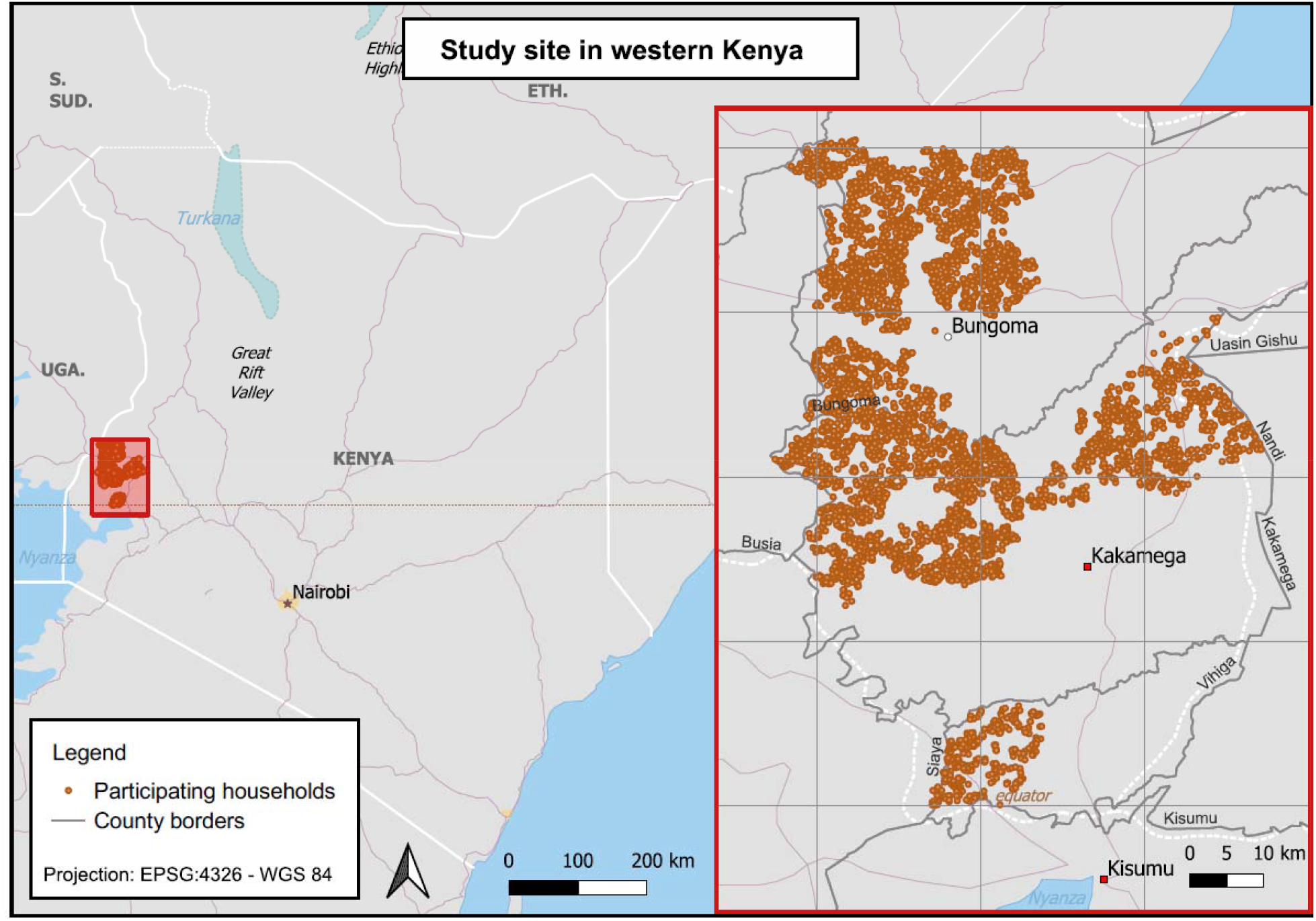
Study Site (2500 km^2^) in western Kenya. Participating households (N=5,761) are plotted.

WASH Benefits visited households prior to intervention delivery (baseline, 2012-2014) and approximately one (midline) and two years (endline) after intervention delivery. Investigators assessed environmental contamination in a subset of households: Study staff collected source water samples (N=1,673) only at baseline; stored water samples from the C/N, WSH/WSHN, W, and H arms at baseline (N=5,761), midline (N=1,577), and endline (N=2,329); and child hand rinse samples from the C/N and WSH/WSHN arms at midline (N=1,026) and endline (N=1,634). Observations from the C and N arms (C/N) and observations from the WSH and WSHN arms (WSH/WSHN) were grouped because nutrition was not expected to impact environmental contamination. GPS coordinates were collected for each water source (baseline) and for each household (baseline, midline, and endline). For stored water collection, study staff asked respondents to show them what they would use if their child 0-3 years old wanted a drink of water. Study staff also sampled the water source that the household reported collecting the stored water from if it was within the same village. All water samples were collected as 150 mL samples in sterile Whirlpak bags. If the respondent reported adding chlorine to the stored water, study staff added sodium thiosulfate to neutralize chlorine residual and measured free chlorine residual using the Hach Pocket Colorimeter II. Child hand rinse samples were collected by filling a Whirlpak bag with 250 mL clean distilled water, placing the index child’s hands in the bag one at a time, massaging the hand, and shaking the hand. More details on this method have been published elsewhere^58–60^. All samples were transported to the field lab on ice and processed the same day of collection. Laboratory technicians analyzed all environmental contamination samples by membrane filtration with MI media (BD, United States) to detect *E. coli* and incubated at 35 degrees Celsius for 20 hours following U.S. Environmental Protection Agency approved method 1604^61^.

We paired gridded meteorological data with point household observations using Google Earth Engine Code Editor, a web-based integrated development environment for the Google Earth Engine JavaScript API. We extracted temperature and precipitation data from the following publicly available gridded sources using the sample collection location GPS coordinates and date of collection:

⍰ *Precipitation:* Climate Hazards Group InfraRed Precipitation with Station data (CHIRPS) is a quasi-global rainfall time series dataset spanning 1981 to present. Daily precipitation is available at 0.05 degree resolution, corresponding to an area approximately 5.6 km by 5.6 km^62^.
⍰ *Temperature:* National Centers for Environmental Prediction (NCEP) Climate Forecast System (CFS) spans 1979 to present. Maximum land surface temperature is available for every 6 hours at 0.2 degree resolution, which corresponds to approximately 22.3 km by 22.3 km^63^.

### Data Analysis

We examined the effects of heavy 7-day precipitation (>90^th^ percentile of our data, 72.7 mm) and high 7-day temperature (mean of daily maximums > 32 degrees C) on environmental *E. coli* levels. We chose threshold predictor variables because we were primarily interested in the effects of extreme weather and did not necessarily expect effects to be linear. This is supported by previous studies, which have observed increased diarrhea risk at heavy rainfall levels^37^. In previous diarrhea risk studies and recent water quality studies, temperature and precipitation have been examined at a variety of timescales (monthly, weekly, daily)^37,40,45,46^. We selected exposures at the weekly level for this analysis based on the assumption that *E. coli* survival would be highest at shorter time scales. We computed 7-day measures using the 7 days prior to sample collection (excluding the day of).

We performed data cleaning and analysis in Stata/MP 16.1 and RStudio version 2022.07.0. For our primary analysis, we used multivariate ordinary least squares (OLS) linear regression to examine the combined effect of heavy precipitation and high temperature on log_10_-transformed *E. coli* levels in source water, stored water, and on child hands. We controlled for WASH Benefits treatment arm in all models.

We considered 8-week precipitation as a potential effect modifier because some studies have found that the association between rainfall and diarrheal illness only holds following prolonged dry periods^35,38,39^. We calculated 8-week precipitation tertiles and repeated the analysis stratified in two subgroups: low precipitation (0^th^ to 33^rd^ percentile) compared to moderate or high precipitation (>33^rd^ percentile). We hypothesized that water treatment may mitigate the effects of weather on water quality. For this reason, we examined self-reported water treatment (any method) and confirmed chlorine water treatment (detectable free chlorine residual) as effect modifiers. We considered improved source versus unimproved source as an effect modifier because we hypothesized that water collected from an unimproved source may be more susceptible to contamination during heavy precipitation events than water collected from an improved source. We also considered specific source type (for source types with > 100 observations) because mechanisms may differ by source type. For example, surface water sources (streams, rivers, lakes, ponds) are generally more open and may be more exposed to sunlight during hot weather (potentially inactivating bacteria) compared with other source types. In addition to conducting stratified analyses, we tested for statistical significance of effect modifiers by including an interaction term in our multivariate models.

A potential weakness of this study is that the results may be sensitive to the chosen predictor variable definitions (7-day period, choice of “heavy/high” thresholds). To mitigate this, we performed several sensitivity analyses with varied specifications. We repeated the analysis using: (1) 5-day time periods rather than 7-day time periods, (2) absolute precipitation and temperature rather than thresholds, (3) the 90^th^ percentile of 7-day mean max temperature (35.8 degrees C) as a threshold for “high temperature”, rather than 32 degrees C, and (4) heavy 1-day precipitation (exceeds 90^th^ percentile, 15.5 mm) during any 1-day period in the previous 7 days rather than total 7-day precipitation.

Because respondents may react to changes in weather, we examined effects of heavy 7-day precipitation (>90^th^ percentile of our data, 72.7 mm) and high 7-day mean maximum temperature (mean of daily maximums > 32 degrees C) on several water-related behavioral measures: whether stored water was collected from an improved source, what source type stored water was collected from, whether the respondent treated the water, what treatment methods were used, and how long the water was stored for. We used multivariate Poisson regression for binary outcomes (improved source, source type, treatment, and treatment method) and multivariate OLS regression for continuous outcomes (water storage time). We controlled for treatment status in all models.

## RESULTS AND DISCUSSION

As previously reported^64^, *E. coli* prevalence was high (>90%) among all water source types and on child hands (Table 1). *E. coli* levels (CFU/100 mL) were lower in household stored water (median = 29 CFU/100 mL) than in water sources (median = 69 CFU/100 mL), potentially because only one sample was collected per source (not accounting for use by multiple households), or because some households (19%) reported treating their water. The median *E. coli* level on child hands was 37 CFU/100 mL.

**Table 1:**
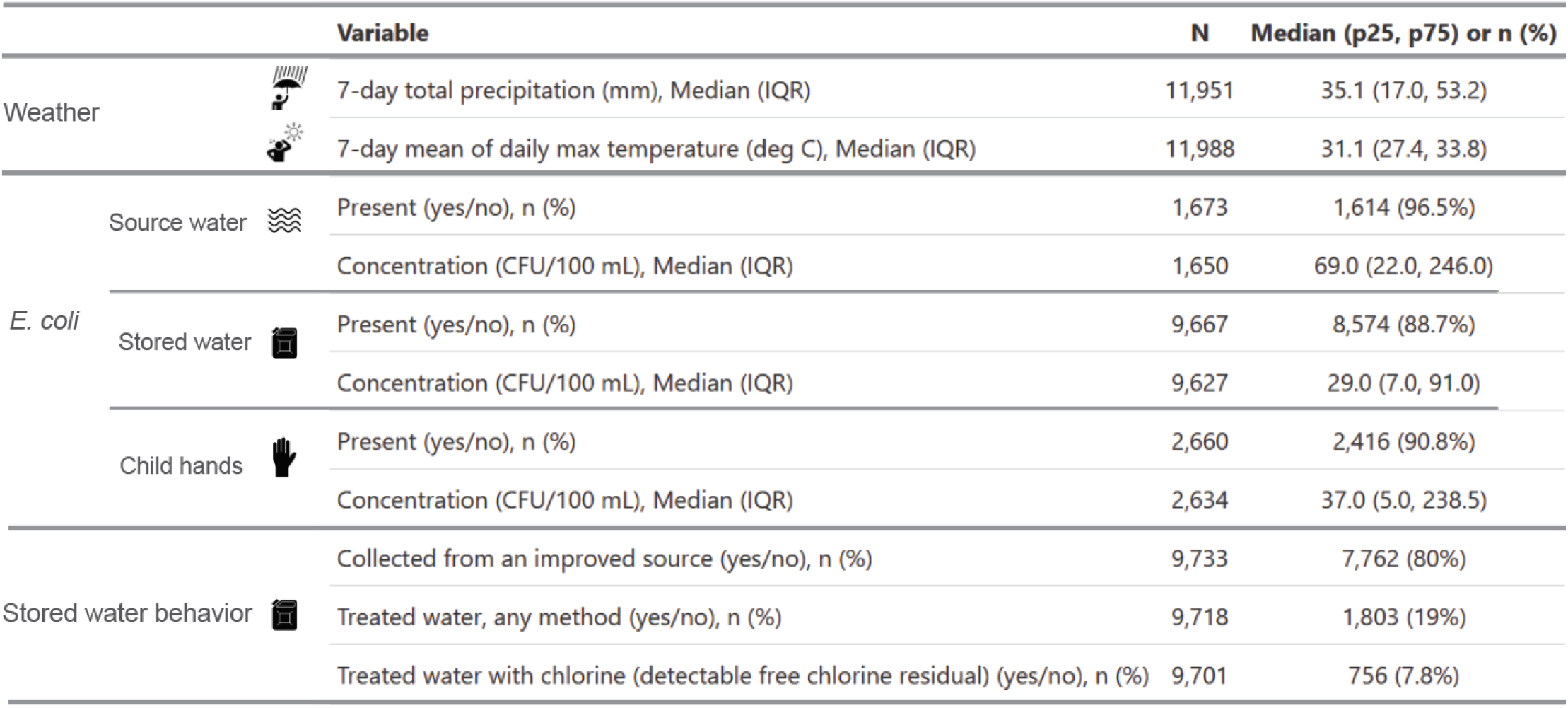
Descriptive Statistics. Sample size (N) is shown for all variables. Median, 25th percentile, and 75th percentile are shown for continuous variables. Prevalence of yes responses (number and percent) is shown for binary variables. Sample sizes are smaller for E. coli concentration variables than for E. coli presence variables because a small number of plates with E. coli colonies were uncountable.

### Primary Analysis

High temperatures and heavy total precipitation during the week before sample collection were significantly associated with environmental *E. coli* levels (Figure 2). In water sources, heavy precipitation (> 90^th^ percentile) was associated with a 0.29 increase in log_10_ CFU *E. coli* per 100 mL water (p < 0.001) and high temperature (mean > 32 degree C) was associated with a 0.16 increase in log_10_ CFU *E. coli* per 100 mL water (p < 0.001). These effects are similar in magnitude to the effect associated with improved sources (reductions of 0.33 log_10_ *E. coli* CFU per 100 mL in water sources and 0.19 log_10_ *E. coli* CFU per 100 mL in household stored water), indicating that the effects of heavy precipitation and high temperature are meaningful. In household stored water, heavy precipitation was associated with a 0.079 increase in log_10_ CFU *E. coli* per 100 mL water (p = 0.042), demonstrating that temperature and precipitation affect water quality at the point that it is being consumed. High temperature was not significantly (p > 0.05) associated with *E. coli* levels in household stored water.

**Figure 2:**
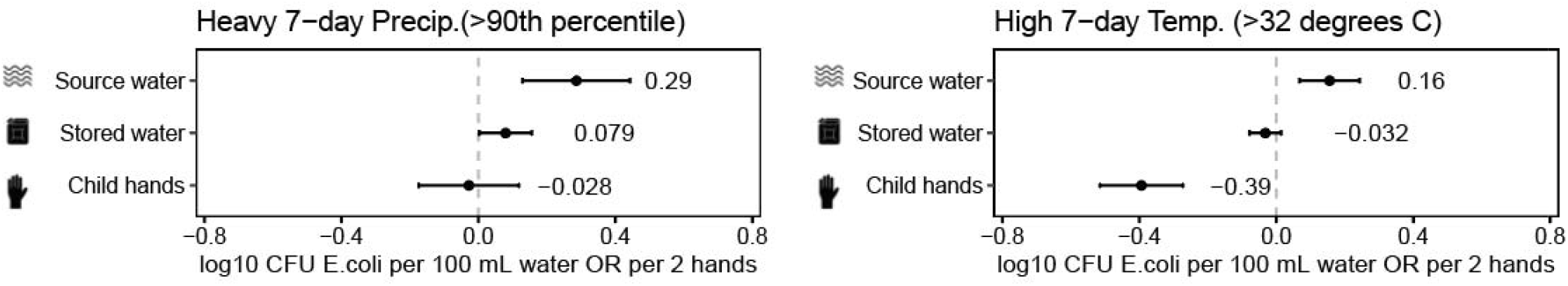
Associations between heavy 7-day precipitation (left), high 7-day temperature (right), and E. coli levels in source water, stored water, and child hands. Point estimates are plotted and labeled. Units are log_10_ CFU E. coli per 100 mL for source water and stored water, and per 2 hands for child hands. Error bars show 95% confidence intervals.

Heavy precipitation was not significantly associated with *E. coli* levels on child hands. However, high temperature was significantly associated with a 0.39 decrease in log_10_ *E. coli* CFU per two hands (p < 0.002), potentially due to faster die-off of bacteria due to heat and sunlight (which is positively correlated with temperature). Temperature is also inversely related to relative humidity, and low relative humidity could reduce bacterial transfer from the household environment (e.g., floor, toys) to child hands. Lopez et al. (2013) found that *E. coli* and most other organisms they examined had lower transfer efficiencies from fomites to fingers at low relative humidity^65^. Temperature is unlikely to influence hand washing effectiveness^66–68^ or hand rinse sample bacterial yield^69^, but it could impact hand washing or bathing frequency because children and their caregivers may use water to cool down when it is hot. During direct observation in north-western Burkina Faso, Traore et al. (2016) found that children swam and mothers bathed under-five children more frequently during a hot period compared with a cold period^70^. Increased bathing or other water contact could have had co-benefits for hand hygiene. Pickering et al. (2011) found that bathing (of self or child) decreased *E. coli* levels on mothers’ hands^71^.

### Effect Modification: 8-Week Rainfall

Low long-term precipitation modified the effect of heavy precipitation on *E. coli* levels in water sources (p = 0.004 on interaction term, Supplementary Table 1), with a larger effect (0.61 log_10_ CFU *E. coli* per 100 mL) after low 8-week rainfall compared to moderate or high 8-week rainfall (Figure 3). Low long-term precipitation also modified the effect of high temperature on *E. coli* levels on child hands (p = 0.045 on interaction term, Supplementary Table 1), with a larger magnitude effect (−0.67 log_10_ CFU *E. coli* per 100 mL) after low 8-week rainfall (Figure 3).

**Figure 3:**
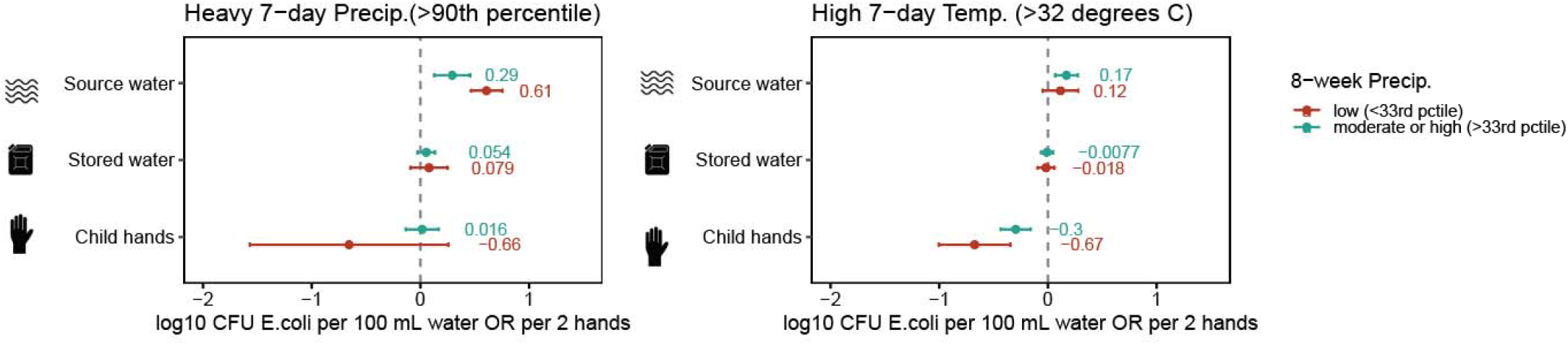
Effect modification by low long-term (8-week) precipitation. Associations between heavy 7-day precipitation (left), high 7-day temperature (right), and E. coli levels in source water, stored water, and child hands. Results are stratified by low (0^th^ to 33^rd^ percentile) vs. moderate or high (>33^rd^ percentile) 8-week rainfall. Point estimates are plotted and labeled. Units are log_10_ CFU E. coli per 100 mL for source water and stored water, and per 2 hands for child hands. Error bars show 95% confidence intervals.

### Effect modification: Water treatment

Water treatment modified the effect of heavy precipitation on *E. coli* levels in stored water (p < 0.001 on interaction term, Supplementary Table 2). While heavy precipitation was associated with increased *E. coli* levels (0.094 log_10_ CFU *E. coli* per 100 mL) in stored water from households who did not treat their water, this relationship did not hold among households who treated their water (Figure 4). Confirmed chlorine water treatment did not modify the effects of heavy precipitation or high temperature on *E. coli* levels in stored water (Supplementary Figure 1, Supplementary Table 3). This analysis was limited by a small number of samples with detectable free chlorine residual (N = 756).

**Figure 4:**
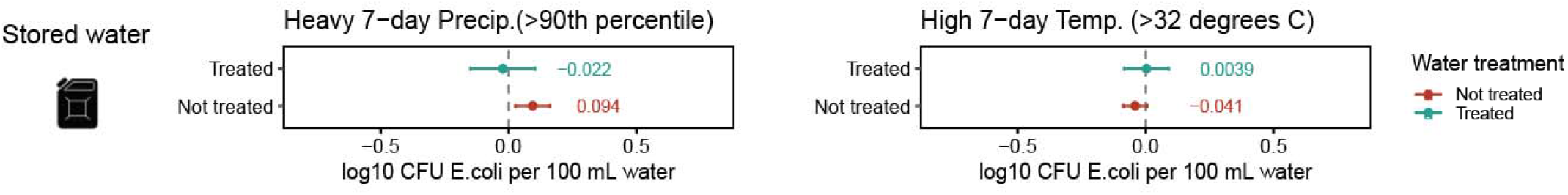
Effect modification by self-reported water treatment. Associations between heavy 7-day precipitation (left), high 7-day temperature (right), and E. coli levels in household stored water. Results are stratified by treated vs. not treated. Point estimates are plotted and labeled. Units are log_10_ CFU E. coli per 100 mL. Error bars show 95% confidence intervals.

### Effect Modification: Source Type

Heavy precipitation was associated with a larger effect on *E. coli* levels in unimproved sources (0.42 log_10_ CFU *E. coli* per 100 mL, p = 0.026 on interaction term, Figure 5A, Supplementary Table 4) compared to water from improved sources, likely because unimproved sources are at higher risk for contamination via runoff or flooding. Heavy precipitation was associated with a larger effect on *E. coli* levels in unprotected springs (0.52 log_10_ CFU *E. coli* per 100 mL, p = 0.005 on interaction term, Figure 5A, Supplementary Table 5) compared with other source types. Unprotected wells versus other source types modified the effect of high temperature on *E. coli* levels in water sources (p = 0.001 on interaction term, Supplementary Table 3) and changed the direction of the effect: Although high temperature increased *E. coli* levels in water sources overall (Figure 2), high temperature reduced *E. coli* levels in unprotected wells (−0.2 log_10_ CFU *E. coli* per 100 mL), potentially because unprotected wells are open and more exposed to sunlight (correlated with temperature) than other source types.

**Figure 5:**
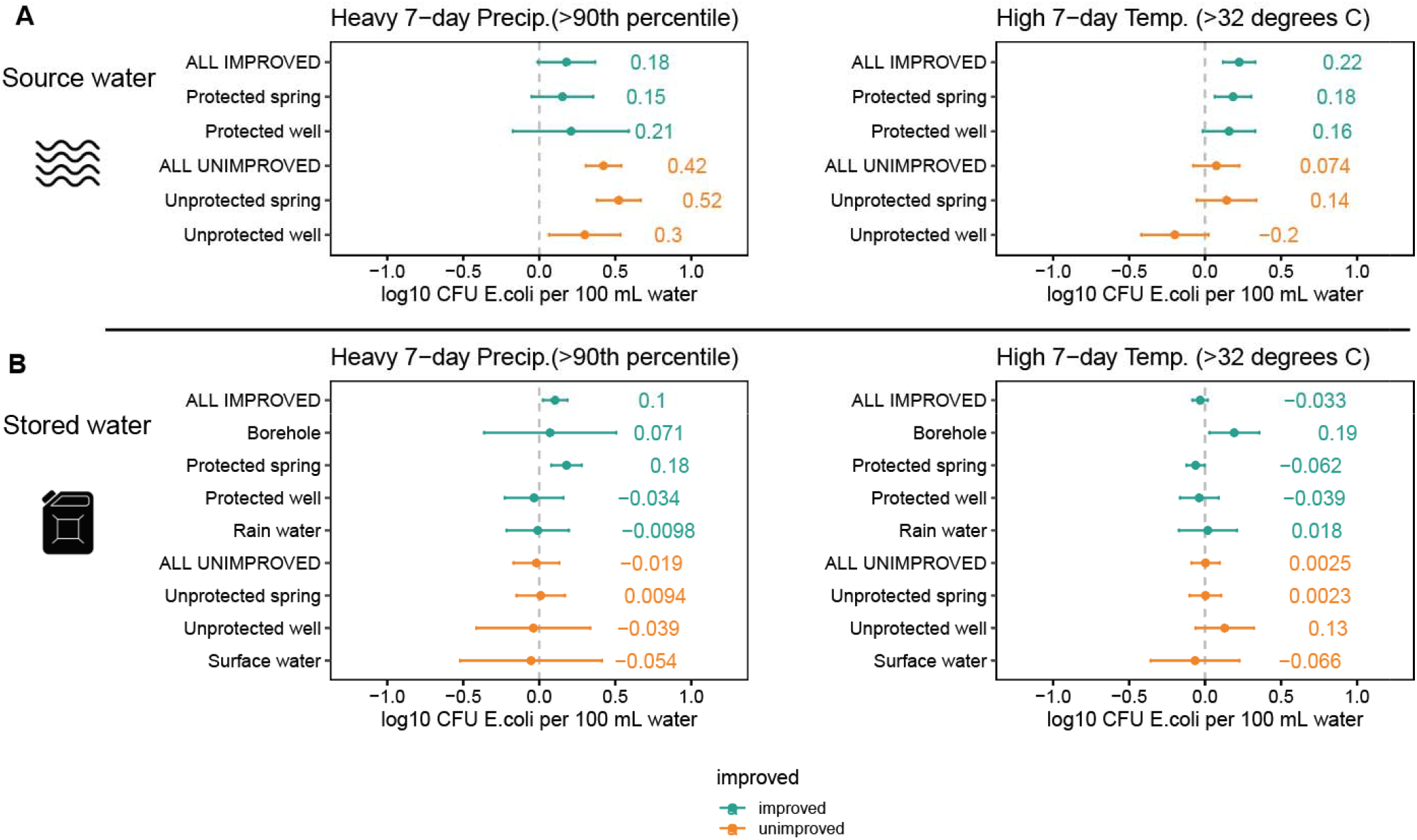
Effect modification by source type. Associations between heavy 7-day precipitation (left), high 7-day temperature (right), and E. coli levels in source water (Figure 5A) and stored water (Figure 5B). For source water, results are stratified by observed source type. For stored water, results are stratified by the source type that the respondent reported collecting the water from. Point estimates are plotted and labeled. Units are log_10_ CFU E. coli per 100 mL. Error bars show 95% confidence intervals.

Source type also modified the effects of heavy precipitation and high temperature on *E. coli* levels in stored water. Heavy precipitation was associated with a larger effect on *E. coli* levels in stored water collected from a protected spring (0.15 log_10_ *E. coli* CFU per 100 mL, p = 0.006 on interaction term, Figure 5B, Supplementary Table 5) compared to water collected from other source types. Collection from boreholes versus other source types modified the effect of high temperature on *E. coli* levels in stored water (p = 0.017, Supplementary Table 5) and changed the direction of the effect: Although high temperature was weakly associated (statistically insignificant, p = 0.26) with decreased *E. coli* levels in stored water overall (Figure 2), high temperature was associated with increased *E. coli* levels in water collected from boreholes (Figure 5B).

### Behavior

Although high temperature was associated with increased *E. coli* levels in water sources, this did not hold true in household stored water (Figure 2), potentially because stored water quality is more complex and impacted by numerous household-level behavioral choices. We found evidence that households altered water-related behaviors (i.e., where to collect water, whether to treat it, how long to store it) in response to the weather. Households were more likely to collect water from an improved source after high 7-day temperature (prevalence ratio = 1.1, p = 0.003, Figure 6B), driven predominantly by an increased likelihood of collection from a protected spring or protected well (improved) and decreased likelihood of collection from an unprotected spring or source water (unimproved) (Figure 6C). This could be due to changes in availability (e.g., some sources may dry up during hot weather) or because respondents choose different water sources based on perceived changes in quality (e.g., color, turbidity, taste). Increased collection from an improved source could mitigate the effect of elevated *E. coli* levels in water sources after high 7-day temperature because collection from an improved source is associated with improved water quality (−0.19 log_10_ CFU *E. coli* per 100 mL in our data). High 7-day temperature was also associated with shorter water storage time (mean difference = 3.81 hours, p = 0.01, Figure 6A), perhaps because households drink^72,73^ or use^74,75^ more water during hot weather. Because water storage time can make water more prone to recontamination^76^, shorter storage time could also mitigate the effects of high temperature.

**Figure 6:**
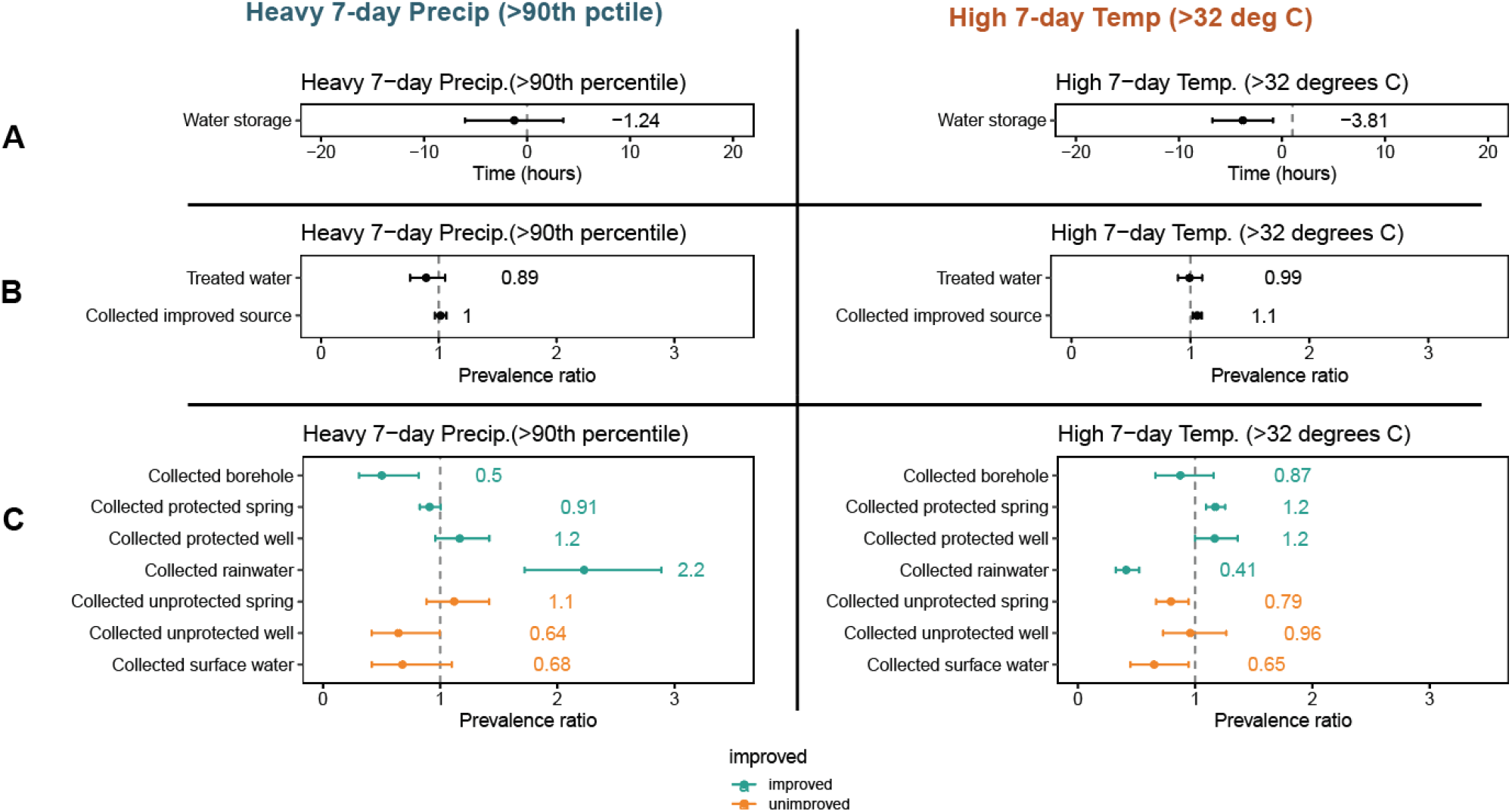
Associations between heavy 7-day precipitation (left), high 7-day temperature (right), and reported stored w ter behaviors: water storage time (Figure 6A), water treatment and collection from an improved source (Figure 6B), and source type that the respondent reported collecting water from (Figure 6C). Water storage time in hours is plotted and labeled for Figure 6A. prevalence ratios are plotted and labeled for Figure 6B and Figure 6C. Error bars in all panels show 95% confidence intervals.

Heavy precipitation increased *E. coli* levels in both water sources and stored water, but the effect magnitude was smaller in household stored water. Heavy precipitation was not significantly associated with the decision to treat water or collect from an improved source (Figure 6B). However households were more likely to collect rainwater after heavy rain (prevalence ratio = 2.2, p <0.001, Figure 6C). Increased rainwater collection (improved source) could diminish the effects of elevated *E. coli* levels in water sources after heavy precipitation because the mechanisms by which heavy precipitation affects water quality in other source types (e.g., runoff) may not apply to rainwater collection (Figure 5). Although rainwater collection was associated with improved water quality in our data (−0.09 log_10_ CFU *E. coli* per 100 mL), roof-harvested rainwater is not always free from microbial^77,78^ and chemical^77,79^ contamination both because contaminants from the atmosphere may be present in rainwater and because contaminants may accumulate on the roof. Respondents were more likely to collect rainwater after heavy 7-day precipitation (prevalence ratio = 2.2, p < 0.001) and less likely to collect rainwater after high 7-day temperature (prevalence ratio = 0.41, p < 0.001) (Figure 6C).

Temperature and precipitation extremes may have also triggered other unobserved behaviora changes that mediate impacts on environmental *E. coli* levels. For example, children may play outside less (potentially reducing exposure) when it has been raining or is very hot. Children and their caregivers may also interact more with water when it is hot out^70^, which could have co-benefits for hand hygiene^71^.

A limitation of this analysis is that data was not collected uniformly over the course of each year. As a result, the available data may not be representative of typical seasonal meteorological conditions for the study area. For example, relatively few source water observations were collected at times of year that typically have low precipitation and high temperatures (January, February) and high precipitation and high temperatures (April, May). However, data collection was spread out such that at least some data was collected in every month of the year.

There are important fecal transmission pathways (e.g., fomites, fields, flies, food) that were beyond the scope of this study. Additional work examining the impact of weather on other pathways would be valuable for anticipating climate change impacts and prioritizing interventions. Because mechanisms and behaviors may vary by context, evidence from additional geographic locations would strengthen our understanding of weather impacts on fecal contamination in water and on hands. Because weather has a significant effect on environmental *E. coli* levels, inclusion of weather variables such as temperature and precipitation may improve the precision of impact evaluations on fecal contamination in the environment even when these are not the primary exposures of interest. Satellite data enables the incorporation of weather data with relative ease.

We found that heavy precipitation and high temperature had meaningful and statistically significant effects on *E. coli* levels in water sources, stored water, and child hands. These effects were consistent across multiple sensitivity analyses (Supplementary Figures 2-5), suggesting that the effects were true and not sensitive to the choice of model specification. In water sources, heavy rainfall increased *E. coli* levels, perhaps by transporting feces via increased runoff and flooding. This finding is consistent with studies in other locations^40,45,46^. The effect was larger after low 8-week rainfall, potentially because long dry periods allowed feces to accumulate in the environment. Heavy rainfall also increased *E. coli* levels in household stored water. Notably, heavy rainfall did not increase *E. coli* levels among the subset of respondents who reported treating their water, suggesting that water treatment can mitigate effects on water quality. In this first study of weather and hand contamination, high temperature decreased *E. coli* levels on child hands. There are a few potential explanations for this finding. Heat and sunlight may have increased *E. coli* die-off. Low relative humidity (inversely related to temperature) may have reduced bacterial transfer from contaminated fomites to child hands^65^. Finally, the effect may have been mediated by behavioral changes, such as increased bathing when it is hot.

Our findings demonstrate that in rural Kenya, extreme weather due to climate change will likely increase bacterial contamination in drinking water but reduce contamination on child hands. We show that individuals react to weather in ways that may mitigate effects on household stored water. We suggest that water treatment is an appropriate mitigation strategy for lessening the health burden of climate change.

## ACKNOWLEDGMENTS

We thank the WASH Benefits study team and participants. This research was financially supported in part by Global Development grant OPPGD759 from the Bill & Melinda Gates Foundation to the University of California, Berkeley, CA, USA, and grant AID-OAA-F-13-00040 from United States Agency for International Development (USAID) to Innovations for Poverty Action. This manuscript was made possible by the generous support of the American people through the USAID. The contents are the responsibility of the authors and do not necessarily reflect the views of USAID or the US Government.

