## Supplemental Information for "Effects of high temperature and heavy precipitation on drinking water quality and child hand contamination levels in rural kenya"

**SUPPLEMENTARY INFORMATION**

SUPPLEMENTARY METHODS

*Treatment arms*

In arms assigned to water treatment intervention, study staff installed chlorine dispensers at public water sources and distributed bottled chlorine to enrolled households. In sanitation arms, study staff installed or upgraded pit latrines with a plastic slab and hole cover, distributed child potties, and distributed scoops for removal of child and animal feces. In handwashing arms, study staff installed two dual dispenser tippy-tap devices per household, one near the latrine and one near the kitchen area. The devices dispensed soapy water (soap was provided) and rinse water via separate pedals. In nutrition arms, study staff distributed lipid-based nutrient supplements. In all intervention groups, promoters visited households at least once every two months.

*Data Cleaning and Verification*

Household GPS coordinates were cleaned by comparing the coordinates collected at baseline, midline, and end line. The baseline latitude was replaced with the average of the midline and end line latitudes if baseline latitude was more than 0.05 degrees different than the midline and end line latitudes, midline and end line latitudes were not missing, and the household had not temporarily or permanently moved at midline or end line. The baseline longitude was replaced with the average of the midline and end line longitudes if baseline longitude was more than 0.05 degrees different than the midline and end line longitudes, midline and end line longitudes were not missing, and the household had not temporarily or permanently moved at midline or end line.

The midline coordinates were set equal to the baseline coordinates unless the household temporarily or permanently moved at midline. The end line coordinates were set equal to the baseline coordinates unless the household permanently moved at midline or temporarily or permanently moved at end line. The coordinates were plotted in ArcMap and visually inspected. Points that were obviously wrong (very far from other points, wrong geographic area) were replaced with the median of the coordinates from households in the same village.

Source water GPS coordinates were cleaned by comparing with the finalized household baseline coordinates. Source coordinates that differed from household coordinates by a total distance of more than 0.07 degrees were replaced with the household coordinate. Source water GPS coordinates were plotted in QGIS and visually examined. No clearly wrong points were identified.

Dates were validated through comparison between the dates recorded in the survey and the date that the samples were reported to be collected. If these dates differed by more than three days, the observations were dropped due to concerns that a failed match could have occurred while merging survey and sample data, and that the date may not be reliable.

Only one sample was considered per water source, even if multiple households reported using this source. Duplicates by sample ID were examined manually. They all appeared to be true duplicates because *E. coli* and enumerator data was identical between observations with the same sample ID. There were differences in the GPS coordinates and household ID, which could be due to the same source being recorded multiple times for different households. Duplicates were dropped.

Study child twins were dropped to avoid duplicates since environmental contamination was observed at the household level, which would be the same for twins.

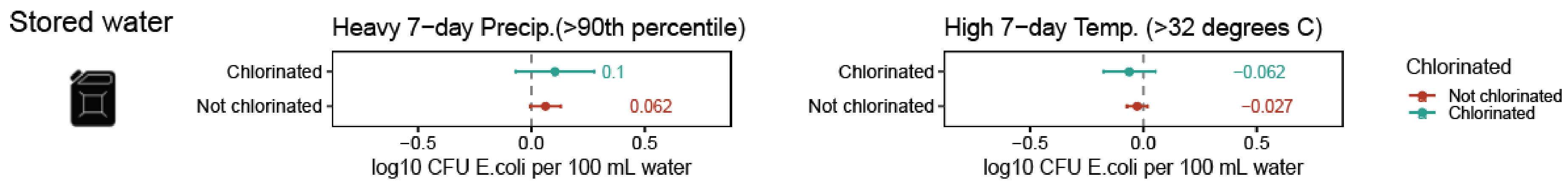


*Supplementary Figure 1: Effect modification by confirmed chlorine water treatment (detectable chlorine residual). Associations between heavy 7-day precipitation (left), high 7-day temperature (right), and E. coli levels in household stored water. Results are stratified by chlorinated vs. not chlorinated. Point estimates are plotted and labeled. Units are $log_{10}$ CFU E. coli per 100 mL. Error bars show 95% confidence intervals.*

*Supplementary Table 1: Statistical significance testing for effect modification by low ($0^{th}$ to $33^{rd}$ percentile) vs. moderate or high (>$33^{rd}$ percentile) 8-week precipitation*

| ***Interaction term*** | ***Sample type*** | ***Coefficient on interaction term ($log_{10}$ CFU E. coli per 100 mL or 2 hands)*** | ***Standard error*** | ***P-value*** | ***Lower 95% CI*** | ***Upper 95% CI*** | ***N*** |
|---|---|---|---|---|---|---|---|
| *7-day high temp and low 8-week precipitation* | *stored* | *-0.076* | *0.052* | *0.147* | *-0.178* | *0.027* | *9261* |
| *7-day heavy precipitation and low 8-week precipitation* | *stored* | *-0.039* | *0.118* | *0.743* | *-0.271* | *0.194* | *9261* |
| *7-day high temp and low 8-week precipitation* | *source* | *-0.055* | *0.096* | *0.567* | *-0.245* | *0.134* | *1613* |
| *7-day heavy precipitation and low 8-week precipitation* | *source* | *0.316* | *0.111* | *0.004* | *0.099* | *0.534* | *1613* |
| *7-day high temp and low 8-week precipitation* | *hands* | *-0.359* | *0.178* | *0.045* | *-0.711* | *-0.007* | *2408* |
| *7-day heavy precipitation and low 8-week precipitation* | *hands* | *-0.673* | *0.511* | *0.190* | *-1.681* | *0.335* | *2408* |

*Supplementary Table 2: Statistical significance testing for effect modification by water treatment vs. no water treatment*

| ***Interaction term*** | ***Sample type*** | ***Coefficient on interaction term (log10 CFU E. coli per 100 mL or 2 hands)*** | ***Standard error*** | ***P-value*** | ***Lower 95% CI*** | ***Upper 95% CI*** | ***N*** |
|---|---|---|---|---|---|---|---|
| *7-day high temp and water treatment* | *stored* | *0.007* | *0.071* | *0.924* | *-0.133* | *0.147* | *9193* |
| *7-day heavy precipitation and water treatment* | *stored* | *-0.324* | *0.090* | *<0.001* | *-0.501* | *-0.148* | *9193* |

*Supplementary Table 3: Statistical significance testing for effect modification by detectable free chlorine residual vs. no detectable free chlorine residual*

| ***Interaction term*** | ***Sample type*** | ***Coefficient on interaction term (log10 CFU E. coli per 100 mL or 2 hands)*** | ***Standard error*** | ***P-value*** | ***Lower 95% CI*** | ***Upper 95% CI*** | ***N*** |
|---|---|---|---|---|---|---|---|
| *7-day high temp and detectable free chlorine residual* | *stored* | *-0.060* | *0.080* | *0.449* | *-0.216* | *0.096* | *9176* |
| *7-day heavy precipitation and detectable free chlorine residual* | *stored* | *-0.073* | *0.105* | *0.487* | *-0.279* | *0.133* | *9176* |

*Supplementary Table 4: Effect modification by improved vs. unimproved source*

| ***Interaction term*** | ***Sample type*** | ***Coefficient (log10 CFU E. coli per 100 mL or 2 hands)*** | ***Standard error*** | ***P-value*** | ***Lower 95% CI*** | ***Upper 95% CI*** | ***N*** |
|---|---|---|---|---|---|---|---|
| *7-day high temperature and improved source* | *stored* | *-0.036* | *0.056* | *0.522* | *-0.145* | *0.074* | *9207* |
| *7-day heavy precipitation and improved source* | *stored* | *0.11* | *0.089* | *0.217* | *-0.065* | *0.284* | *9207* |
| *7-day high temperature and improved source* | *source* | *0.151* | *0.091* | *0.098* | *-0.028* | *0.329* | *1613* |
| *7-day heavy precipitation and improved source* | *source* | *-0.244* | *0.109* | *0.026* | *-0.458* | *-0.03* | *1613* |

*Supplementary Table 5: Statistical significance testing for effect modification by source type (each source type vs. all others)*

| ***Interaction term*** | ***Sample type*** | ***Coefficient on interaction term (log$_{10}$ CFU E. coli per 100 mL or 2 hands)*** | ***Standard error*** | ***P-value*** | ***Lower 95% CI*** | ***Upper 95% CI*** | ***N*** |
|---|---|---|---|---|---|---|---|
| *7-day heavy precipitation and protected spring* | *source* | *-0.223* | *0.134* | *0.096* | *-0.485* | *0.040* | *1613* |
| *7-day heavy precipitation and protected well* | *source* | *-0.101* | *0.208* | *0.627* | *-0.511* | *0.308* | *1613* |
| *7-day heavy precipitation and unprotected spring* | *source* | *0.338* | *0.122* | *0.006* | *0.098* | *0.578* | *1613* |
| *7-day heavy precipitation and unprotected well* | *source* | *0.024* | *0.143* | *0.870* | *-0.258* | *0.305* | *1613* |
| *7-day high temperature and protected spring* | *source* | *0.044* | *0.079* | *0.575* | *-0.111* | *0.199* | *1613* |
| *7-day high temperature and protected well* | *source* | *0.020* | *0.097* | *0.837* | *-0.170* | *0.210* | *1613* |
| *7-day high temperature and unprotected spring* | *source* | *-0.030* | *0.107* | *0.780* | *-0.241* | *0.181* | *1613* |
| *7-day high temperature and unprotected well* | *source* | *-0.384* | *0.117* | *0.001* | *-0.614* | *-0.154* | *1613* |
| *7-day heavy precipitation and borehole* | *stored* | *-0.174* | *0.221* | *0.430* | *-0.607* | *0.259* | *9210* |
| *7-day heavy precipitation and piped* | *stored* | *0.082* | *0.278* | *0.769* | *-0.463* | *0.627* | *9210* |
| *7-day heavy precipitation and protected spring* | *stored* | *0.195* | *0.071* | *0.006* | *0.056* | *0.335* | *9210* |
| *7-day heavy precipitation and protected well* | *stored* | *-0.131* | *0.106* | *0.216* | *-0.338* | *0.076* | *9210* |
| *7-day heavy precipitation and rain* | *stored* | *-0.112* | *0.118* | *0.342* | *-0.343* | *0.119* | *9210* |
| *7-day heavy precipitation and surface water* | *stored* | *-0.107* | *0.262* | *0.682* | *-0.621* | *0.406* | *9210* |
| *7-day heavy precipitation and unprotected spring* | *stored* | *-0.086* | *0.097* | *0.375* | *-0.275* | *0.104* | *9210* |
| *7-day heavy precipitation and unprotected well* | *stored* | *-0.131* | *0.186* | *0.481* | *-0.497* | *0.234* | *9210* |
| *7-day high temperature and borehole* | *stored* | *0.211* | *0.088* | *0.017* | *0.038* | *0.384* | *9210* |
| *7-day high temperature and piped* | *stored* | *-0.035* | *0.207* | *0.865* | *-0.441* | *0.370* | *9210* |
| *7-day high temperature and protected spring* | *stored* | *-0.059* | *0.048* | *0.219* | *-0.153* | *0.035* | *9210* |
| *7-day high temperature and protected well* | *stored* | *-0.027* | *0.072* | *0.714* | *-0.169* | *0.116* | *9210* |

| | | | | | | | | |
|---|---|---|---|---|---|---|---|---|
| *7-day high temperature and rain* | *stored* | *0.093* | | *0.098* | *0.340* | *-0.099* | *0.285* | *9210* |
| *7-day high temperature and surface water* | *stored* | *0.007* | | *0.161* | *0.964* | *-0.309* | *0.323* | *9210* |
| *7-day high temperature and unprotected spring* | *stored* | *0.025* | | *0.062* | *0.681* | *-0.096* | *0.147* | *9210* |
| *7-day high temperature and unprotected well* | *stored* | *0.143* | | *0.102* | *0.162* | *-0.058* | *0.343* | *9210* |

*Sensitivity*

Results were largely consistent across choices of model specification.

The effects of 5-day weather were similar to that of 7-day weather (Figure 2, Supplementary Figure 2). As with 7-day time periods, heavy 5-day precipitation (>90$^{th}$ percentile) and high 5-day temperature (>32 degrees C) were associated with increased *E. coli* levels in source water. Although 7-day heavy precipitation was significantly associated with *E. coli* levels in household stored water, 5-day heavy precipitation was not significantly associated with *E. coli* levels in household stored water (Supplementary Table 2). However, the direction of the effect was consistent. Effects on child hands were consistent: high temperature was associated with decreased log10 *E. coli* levels and heavy precipitation was not significantly associated with *E. coli* levels.

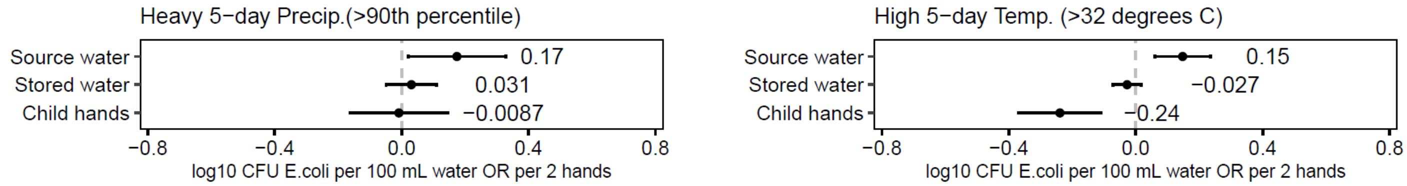


*Supplementary Figure 2: Sensitivity analysis using 5-day periods rather than 7-day periods. Associations between heavy 7-day precipitation (left), high 7-day temperature (right), and E. coli levels in source water, stored water, and child hands. Point estimates are plotted and labeled. Units are log10 CFU E. coli per 100 mL for source water and stored water, and 2 hands for child hands. Error bars show 95% confidence intervals.*

The use of absolute temperature and precipitation rather than thresholds yielded consistent results across all source types (Figure 2, Supplementary Figure 3).

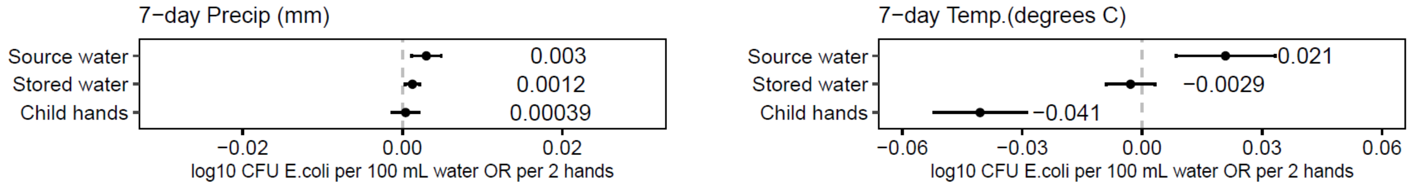


*Supplementary Figure 3: Sensitivity analysis using absolute precipitation and temperature rather than thresholds. Associations between heavy 7-day precipitation (left), high 7-day temperature (right), and E. coli levels in source water, stored water, and child hands. Point estimates are plotted and labeled. Units are log10 CFU E. coli per 100 mL for source water and stored water, and 2 hands for child hands. Error bars show 95% confidence intervals.*

Defining high 7-day mean max temperature using the 90$^{th}$ percentile (35.8 degrees C) rather than 32 degrees C yielded consistent results across all source types (Figure 2, Supplementary Figure 4).

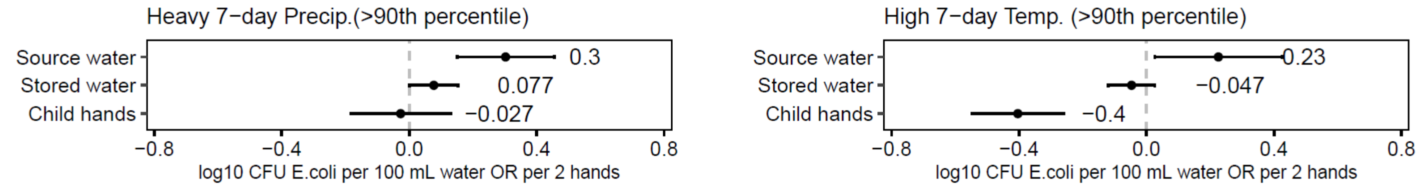


*Supplementary Figure 4: Sensitivity analysis using the 90th percentile 7-day mean maximum temperature (35.8 deg C) rather than 32 deg C. Associations between heavy 7-day precipitation (left), high 7-day temperature (right), and E. coli levels in source water, stored water, and child hands. Point estimates are plotted and labeled. Units are log10 CFU E. coli per 100 mL for source water and stored water, and 2 hands for child hands. Error bars show 95% confidence intervals.*

Considering heavy rainfall events (1 or more days exceeds the 90th percentile in any of the previous 7 days) rather than 7-day total rainfall yielded slightly different results: effects on source water were no longer statistically significant (Figure 2, Supplementary Figure 5). However, effects on stored water were consistent and the direction of both effects were consistent.

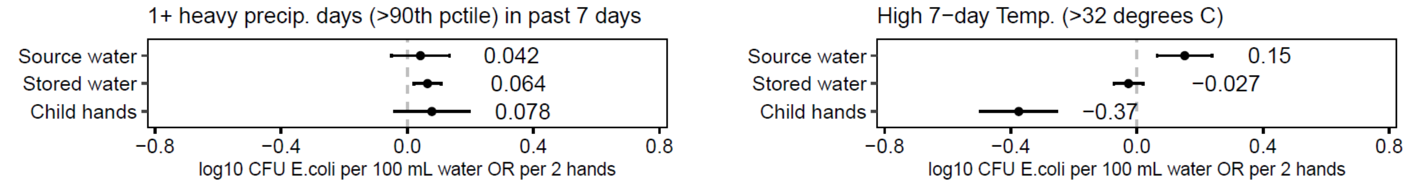


*Supplementary Figure 5: Sensitivity analysis using heavy rainfall events (1 or more days exceeds the 90th percentile in any of the previous 7 days) rather than 7-day total rainfall. Associations between heavy 7-day precipitation (left), high 7-day temperature (right), and E. coli levels in source water, stored water, and child hands. Point estimates are plotted and labeled. Units are log10 CFU E. coli per 100 mL for source water and stored water, and 2 hands for child hands. Error bars show 95% confidence intervals.*
